# Predicting Causal Relationships from Biological Data: Applying Automated Casual Discovery on Mass Cytometry Data of Human Immune Cells

**DOI:** 10.1101/122572

**Authors:** Sofia Triantafillou, Vincenzo Lagani, Christina Heinze-Deml, Angelika Schmidt, Jesper Tegner, Ioannis Tsamardinos

## Abstract

Learning the causal relationships that define a molecular system allows us to predict how the system will respond to different interventions. Distinguishing causality from mere association typically requires randomized experiments. Methods for automated causal discovery from limited experiments exist, but have so far rarely been tested in systems biology applications. In this work, we apply state-of-the art causal discovery methods on a large collection of public mass cytometry data sets, measuring intra-cellular signaling proteins of the human immune system and their response to several perturbations. We show how different experimental conditions can be used to facilitate causal discovery, and apply two fundamental methods that produce context-specific causal predictions. Causal predictions were reproducible across independent data sets from two different studies, but often disagree with the KEGG pathway databases. Within this context, we discuss the caveats we need to overcome for automated causal discovery to become a part of the routine data analysis in systems biology.

## Introduction

Inferring causal relationships is of paramount importance in molecular biology. Knowing the causal structure of a molecular process allows advanced reasoning on its behavior, and such knowledge could be valuable in therapeutic approaches, for example though predicting side effects of pharmaceutical drugs. Given two factors A and B that affect each other, causal knowledge confers more information as compared to correlation. The correlation of factor A and factor B allows to predict the levels of one given the levels of the other however, it gives no information on the possible change in A if B is perturbed. This is not true for causal knowledge: Knowing the causal relationship of the two compounds allows predicting their response to an external intervention. For example, if A causally affects the abundance of B, then the perturbation of A is expected to affect the levels of B, while the perturbation of B will have no impact on A.

Computational causality has developed a language to describe, quantify and reason with causal claims. The most common framework of computational causality is causal Bayesian networks (CBNs), that use a simple assumption to connect causal relationships to associative patterns^1^. CBNs use directed acyclic causal graphs to describe the causal relationships and connect them to associations expected to hold or vanish in the joint probability distribution. Causal effects can also be computed using CBNs using do-calculus, a formal system for causal reasoning that includes an operation for interventions^1^. Algorithms for automatically identifying CBNs from limited or without experiments have also been proposed^2^.

The typical approach to learning causal relationships in current biology is by carrying out specifically designed experiments. Links that are established in the literature are then manually synthesized into larger causal models, like for example the Kyoto Encyclopedia of Genes and Genomes (KEGG) pathway database^3^. Lately, high throughput techniques such as mass cytometry or single cell sequencing allow multivariate interrogation of large numbers of cells under a plethora of experimental conditions, with advanced technical reliability. Thus, applying techniques from the field of computational causality to systems biology could help revolutionize the time consuming, expensive and error-prone process of experimentally identifying causal structure in molecular systems.

In a seminal paper for applied causal discovery^4^, the authors were able to almost flawlessly reconstruct a known causal signaling pathway from a mixture of experimental and observational flow cytometry measurements. This work illustrates the feasibility of causal discovery in biology. However, we must point out that several factors aided this success: the set of variables included in the analysis were not affected by any known latent confounders, and a mixture of observations and perturbations were included, facilitating the correct orientation of the recovered edges. The known system also included a limited number of feedback cycles (which were not identified correctly by the algorithm). De novo identification of causal relationships in data where latent confounders and feedback loops are possible, and interventions do not necessarily have known target, can be far more challenging. A major contribution of this work is to elucidate the challenges of de novo identification of causal relationships.

In this work, we attempt to discover novel causal relationships from a large collection of public mass cytometry data of immune cells perturbed with a variety of compounds. Like flow cytometry, mass cytometry is a technique that can be used to singularize cells and measure protein abundance on the cellular level, resulting in very large sample sizes that are suitable for causal discovery methods. We discuss how different types of experiments can be modeled in the context of causal discovery, and then test the applicability of two state-of-the-art methods to identify phosphorylation of signaling proteins among the measured variables. We test the reproducibility of algorithmic results in similar, albeit different, data sets. Finally, we examine whether algorithmic findings are validated in the literature and in experimental data from the same study.

We find that (a) results are highly consistent on data sets that include different donors, experimental cell-stimulation time-points, or cell types (b) different causal methods often disagree with each other and with known pathways, (c) validity in experimental data is inconclusive. Our approach also revealed a large variation of the correlation structure, even in technical replicates produced within the same lab. MATLAB code for reproducing the results is available in https://github.com/mensxmachina/Mass-Cytometry. These results indicate that (a) de novo discovery of causal pathway relations is still a challenging task for current causal discovery methods, despite the positive results in^4^ and (b) current causal discovery methods do identify reproducible findings across similar data sets. There results constitute an important step for further developments in order to apply causal discovery methods successfully to biological single cell data in the future.

## Results

### Mass cytometry data

As a test bed dataset to assess the applicability and performance of automated causal discovery methods, we used a collection of mass cytometry data published in^5^ (denoted ‘BDM’). Multiplexed mass cytometry was used to measure the abundance of surface proteins and intracellular phosphorylated proteins in human peripheral blood mononuclear cells (PBMCs) that were stimulated with different factors (denoted ‘activators’) in vitro. The measured surface molecules are typically associated with certain PBMC subpopulations of specific function which, thus, can be distinguished by these markers. Within these cell populations, intracellular proteins specifically respond to certain extracellular signals to initiate regulatory pathways. To test these responses, known post-translational phosphorylations of specific proteins involved in immune cell signaling pathways were measured before and 30 minutes after treatment with several suitable activators.

The BDM study uses 11 different activators to stimulate cells, and furthermore it includes a multiple inhibitor experiment measuring protein responses in activated and non-activated cells under 8 dosages of 27 different inhibitors. For each inhibitor, the lower dosage is practically considered equal to zero, resulting in a collection of 27 replicates for the dose “zero”. These data sets serve as a unique test bed for the reproducibility and consistency of algorithmic results. The measurement of the same system under slightly different experimental conditions should reflect similar mechanisms. However, it is worth noting that this is one of the first mass cytometry studies, and variation in the results could in part be attributed to lack of custom normalization techniques developed after its publication^6^.

In the original publication, PBMC samples were manually gated into subpopulations based on the abundance of surface markers. The analysis is performed on this gated data, so each prediction is subpopulation-specific. Figure 1 illustrates the available data sets for each single subpopulation. Different subpopulations have different roles in the immune system and will have different responses to external cues. Moreover, the abundance of proteins within a cell greatly depends on cell size. Pooling data from different subpopulations together would create spurious relationships confounded by cell size. However, we assume that similar subpopulations should exhibit similar behavior.

**Figure 1.**
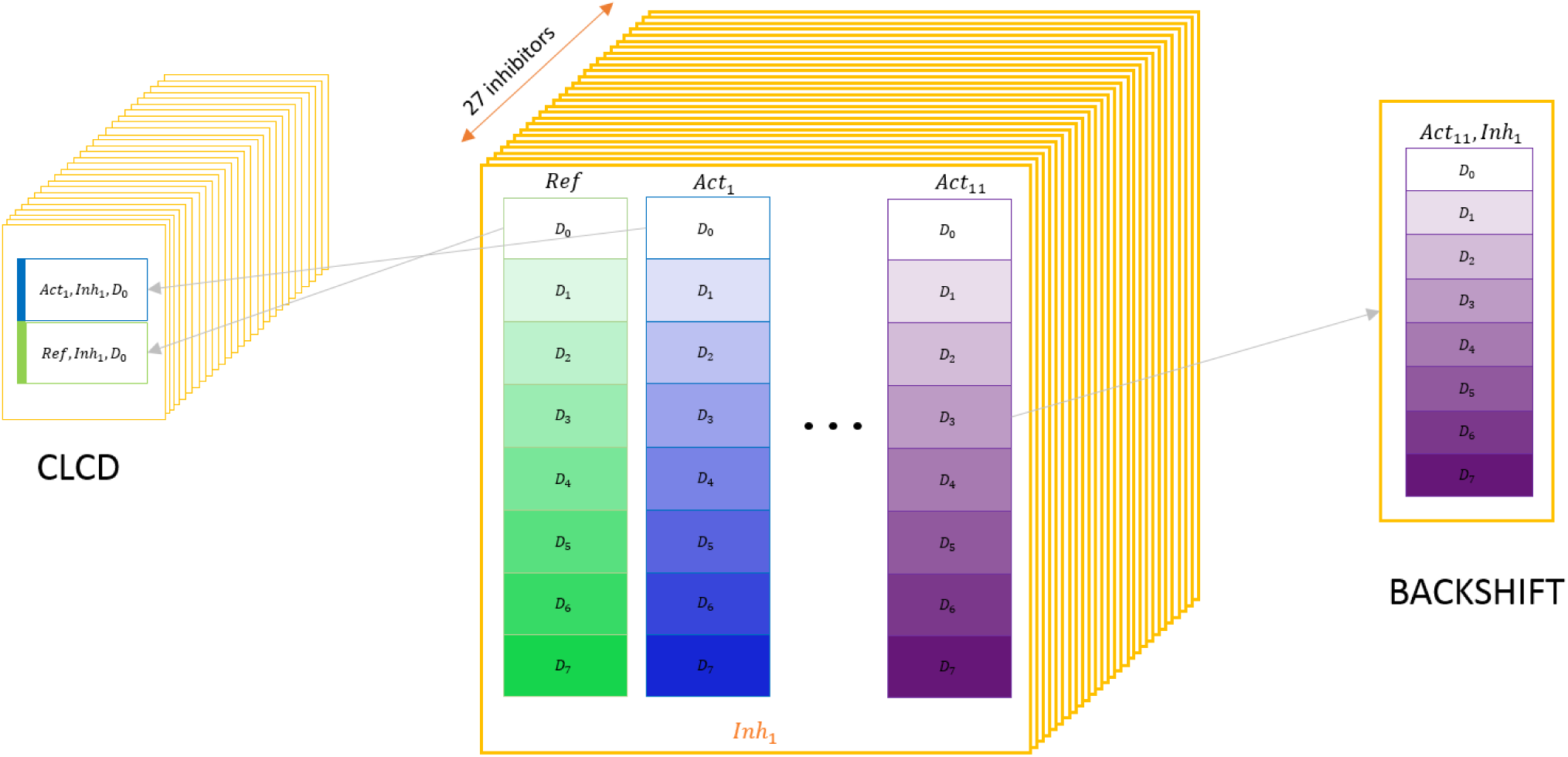
Collection of public mass cytometry data sets used for causal discovery. (middle) Available data for each subpopulation: single cells from a healthy human donor were selected and treated with 8 different dosages of a single inhibitor (*D*_0_ corresponding to the zero dosage), and then stimulated with 11 different activators or without activator (denoted ‘reference’ data with no activation). The experimental plate setup with the different activations was repeated for 27 different inhibitors (with 8 doasages each). (left) Data usage for one run of CLCD: only zero dosages were used. Activated and non-activated data with no inhibition were pooled together to form a new data set which includes a binary column indicating activation status. The process was repeated for all 27 plates (inhibitors), and predictions were decided based on all 27 data sets. (right) Data usage for one run of BackShift: for every activator and inhibitor, different dosages of the same inhibitor are used as different environments.

Surface markers were used to discriminate subpopulation types and are not expected to react to the activator conditions, particularly within a narrow time frame like the one used in this study. We therefore only included functional proteins (measured as phosphorylated proteins whose phosphorylation status can change rapidly as response to activation) in all subsequent analyses.

### Learning causal relations using conditional independence tests

Causal discovery makes assumptions on the nature of causality that connect the observable data properties (i.e., the joint probability distribution of the observed variables) to the underlying causal structure. The most popular causal assumption, famous for inspiring Bayesian networks, is the Causal Markov (CM) assumption. The CM assumption states that every variable is independent of its non-effects given its direct causes.

In the context of computational causality, the causal structure of a set of variables is often modeled using directed graphs. A directed edge from X to Y denotes that X causes Y *directly* in the context of measured variables; no variable included in the model mediates the relationship.

Direct causes and non-effects of a variable can now be identified from the causal graph. The CM assumption connects a given causal graph with a set of conditional independencies (CIs). For example, the causal model shown in Fig. 2e1 implies that in the joint probability distribution of A, S and T, A must be independent of T given S (symb. A ╨ T | S), while all variables are pairwisely dependent. Essentially, this conditional independence denotes that the conditioning variable (S) *mediates* the relationship of the variables it renders independent: any information A carries about T goes through S. Therefore, given the value of S, A and T share no additional information about each other.

**Figure 2.**
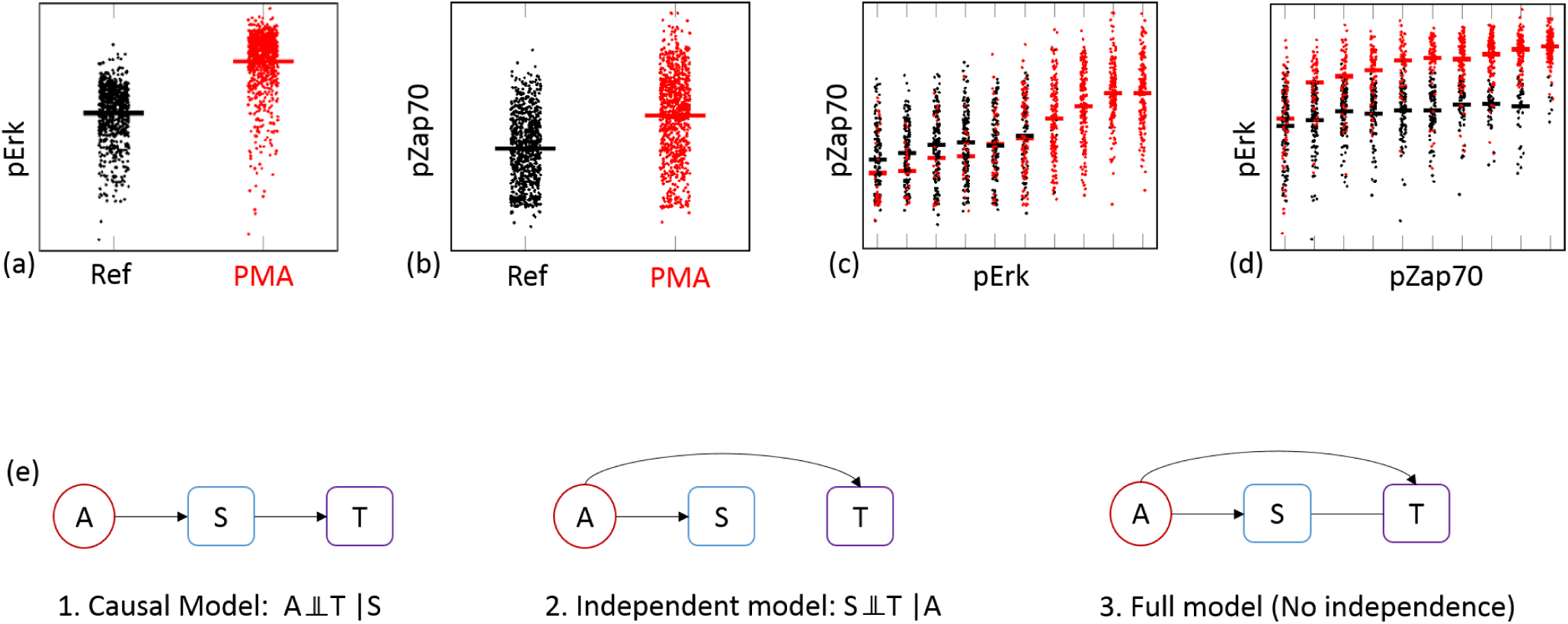
Causal discovery using conditional independence. (a-d) Joint and conditional distributions under conditional independence. Black/red points indicate no activation/activation with PMA. Black/red bars show the mean values. Conditional protein is discretized in10 levels. (a, b) Stimulation with PMA significantly increases the levels of both pErk and pZap70. (c) pZap70 is independent of PMA stimulation given the levels of pErk: P(pZap70|PMA, pErk) = P(pZap70|pErk); mean values for red and black points are very similar. (d)In contrast, PMA stimulation is associated with pErk even after conditioning on pZap70. (e) Possible causal Bayes networks for three variables that are pairwisely dependent, and one of them is restricted to be a source variable (no incoming edges): conditional independence can distinguish among the causal, independent and full model. In the full model, the line between variables S and T does not have any arrowheads; causal direction of this edge cannot be determined from the data.

Figure 2a-d shows an example of a conditional independence pattern in mass cytometry data. Treating natural killer cells with PMA induces the phosphorylation levels of protein Zap70 (pZap70). Thus, stimulation with PMA and pZap70 are dependent: the probability distribution of pZap70 values given PMA stimulation is different than the marginal probability of pZap70 (Fig. 2a). However, given the levels of phosphorylation of protein Erk (pErk), stimulation with PMA is no longer informative for pZap70 (Fig. 2c; small differences in PMA-induced vs Reference pZap70 that are evident for small values of pErk are not statistically significant).

Conditional independencies can be tested in the data using appropriate tests of independence. Computational causal discovery tries to reverse engineer the causal graph using tests of independence: given a pattern of conditional independencies, a causal discovery algorithm typically tries to identify the causal graph that is associated to this pattern through the CM assumption. To do so efficiently, algorithms assume that all observed independencies in the data are a product of the causal structure, rather than being “accidental” or “fine-tuned” properties of the model parameters. This assumption is known as the Faithfulness assumption^2^, and it is employed by most causal discovery algorithms. Even though random parameters typically lead to violations of faithfulness with probability zero^7^, its validity has been debated^8–10^.

Most of the times, CM and faithfulness do not uniquely associate a CI pattern with a single causal model. For example, all causal graphs in Supplementary Fig. 1 are associated with the (single) conditional independence A ╨ T | S. However, additional background knowledge can be used to narrow the space of possible models and enable novel causal discoveries. For example, knowing that A has no causes among the measured variables would rule out all other models apart but the causal model in Fig. 2e1, and allow a novel causal prediction: S must be a cause of T. This simple structure has been successfully exploited in genome-wide association studies (GWAS), where Mendelian randomization is used as a variable that can not be caused by any variable measured in the same study^11^.

This network of three variables, one of which is restricted to be a source node in the causal network, is probably the minimal example where a conditional independence pattern indicates a non-trivial causal relationship (i.e. the mediator is a cause of the third non-source variable). One can identify such CI patterns in large data sets, provided one of the variables is known to have no causes within the model.

Compared to trying to reconstruct the entire network of measured variables, considering only a small subset of variables at each run has many advantages for de novo causal discovery. Causal discovery algorithms are notoriously prone to error propagation^12^, and a single error in a conditional independence test can affect seemingly remote parts of the learnt network. Employing a local approach allows testing all possible conditional independencies, and minimizes the probability that an independence test of unrelated variables will affect the output.

Moreover, limiting the methods to just a few variables reduces the risk of assumption violations. Notice that, in the possible networks shown in Fig. 2, possible latent confounders and/or feedback cycles are not taken into consideration. However, the presence of latent confounders does not affect the validity of the local causal discovery rule presented above: the causal model shown in Fig. 2 is the only model that implies the conditional independence of A and T given S, for all possible networks, even in the presence of confounders (see Supplementary Fig. 2). Feedback cycles are also relatively less troubling, since the only plausible feedback is a loop from S to T (excluding self-loops).

In mass cytometry experiments, the presence of an activator can be modeled as an external, binary variable, that is set by the experimenter and is not influenced by protein phosphorylation within the cell. All causal models where the activator is caused by a phosphorylated protein can then be excluded, and the conditional independence can only be explained if the mediation of S is causal. For example, the conditional independence of PMA and pZap70 given pErk indicates that the correlation between pErk and pZap70 is a causal signal.

Notice that the data in Fig. 2a-d show the levels of pErk and pZap70 with and without PMA stimulation for the zero dosage of the Syki inhibitor. Similar data from the zero dosages of the remaining 26 inhibitors should agree with this CI pattern. Differences in the CI patterns in the 27 conceptually identical data sets could be accounted for by: (1) errors in the results of the conditional independence tests (2) effective inhibition even from a minuscule inhibitor dosage resulting in a different joint probability distribution.

Errors in the results of conditional independence tests can be either Type I errors (reject independence while it holds) or Type II errors (failing to reject an independence that does not hold). Type I errors can often be attributed to measurement noise, as measurement error in the mediating protein can result in failing to identify the conditional independence. Type II errors can be a result of weak dependencies and low sample sizes. In this work, we use two thresholds: a 0.001 threshold for rejecting independence, and a 0.15 threshold for accepting independence. This approach is more conservative than using a single threshold for accepting or rejecting independence. We also consider only triplets for which the same conditional independence pattern holds in at least 10 out of the 27 replicate data sets. Input data for this conservative local causal discovery (CLCD) pipeline are assembled from the original set of data as shown in Fig. 1.

### Learning causal relationships using inhibitors as interventions with unknown targets

CLCD only uses a small portion of the available data, as it disregards all but the lowest inhibitor dosage. Data from different dosages of the same inhibitor correspond to different distributions and cannot be pooled together for CLCD. While some methods exist for integratively analyzing data from multiple experiments^13–15^, the experiments are often required to be surgical interventions (meaning that the targets of the experiments must be known and their values completely set by the experimental conditions). In the available data, different inhibitors can have various unknown targets, while different dosages of the same inhibitor change the magnitude of the effects.

In the recently developed method BackShift ^16^, different experimental conditions (or environments) with unknown targets are used to uncover causal structure. Different experimental conditions are modeled as so-called “shift interventions”, meaning that the values of unknown targets of each intervention are shifted by realizations from a random variable modeling this intervention. Thus, the experiments are not required to be surgical interventions. In contrast, the underlying causal structure is assumed to persist in the experimental condition, affecting the system in addition to the induced intervention effect. The method then exploits the presence of different environments to identify the causal structure of the measured variables, as well as the location and strength of the interventions in each experiment.

To this end, the method assumes a linear causal model that possibly includes cycles and confounders. An example (without cycles nor confounders) is shown in Fig. 3. The coefficients *b_ij_* fully describe the causal structure of the measured variables. Non-zero coefficients of the linear system correspond to direct causal relationships and their value corresponds to their strength. In Fig. 3, all *b_ij_* coefficients apart from *b_xy_* and *b_yz_* are zero.

**Figure 3.**
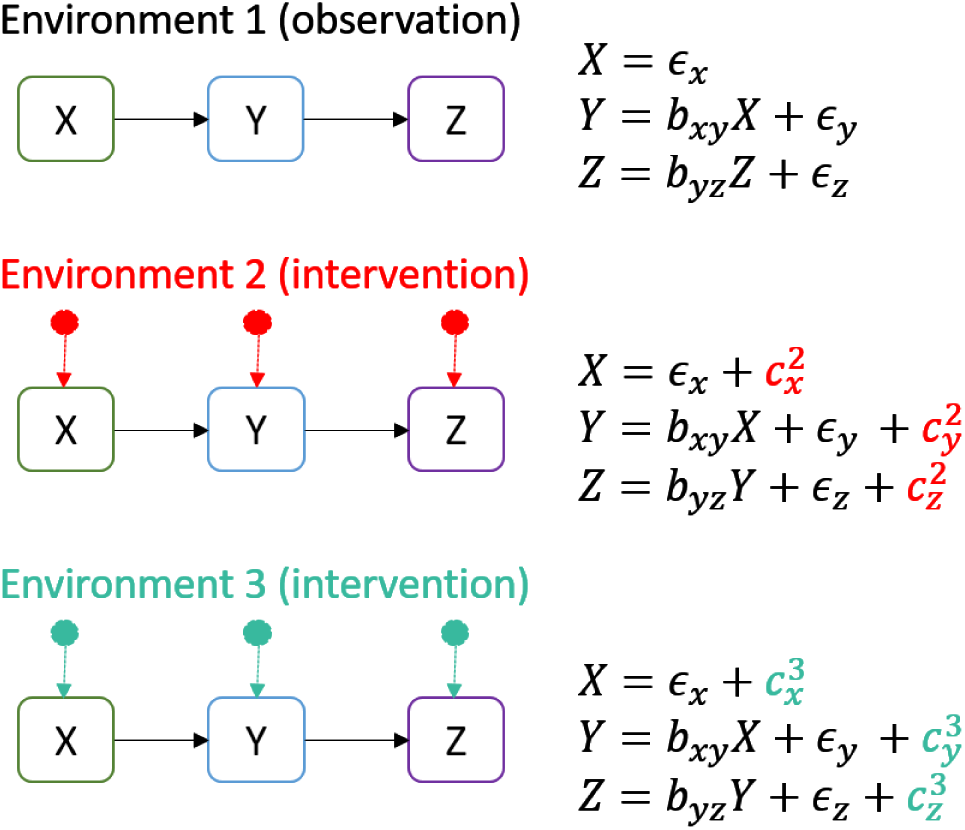
Causal discovery using different experimental conditions. A linear causal model is used to describe the causal structure of the measured variables. Different experimental conditions (environments) possibly “shift” the values of some of the variables (i.e.*c^i^_j_* can be zero for some *i, j*) while the causal structure of the variables remains invariant. BackShift exploits this invariance and can identify the causal structure given observations from at least three different environments.

An intervention is modeled by the random variable *c^i^_j_*, where *i* denotes the experimental condition and *j* denotes the variable the intervention is acting on. The targets of the intervention do not have to be known and we do not require interventions to occur at every variable in each experimental conditions. In other words, some of the *c^i^_j_* can be zero.

The distribution of the noise terms as well as the coefficients *b_i__j_* are assumed to remain invariant across different experimental conditions. The estimation of *b_i__j_* then relies on a simple joint matrix factorization. At least three different environments, one of which can contain purely observational data, are required for identifiability. In this work, different dosages of the same inhibitor were used as different environments (i.e. dosages *D*_0_ to *D*_7_ shown in Fig. 1).

In the presence of various model violations, the joint diagonalization BackShift relies on is not possible. Therefore, model violations can be detected by the success or failure of the joint diagonalization algorithm. Inter alia, this applies to the following situations: (i) an intervention acted on hidden variables; (ii) interventions in the same environment are correlated; (iii) interventions act on the children of a certain variable which has more than one child. To increase robustness of the predictions, we used stability selection^17^ which provides a finite sample control on the number of false discoveries.

As different inhibitor/activator combinations may behave in different ways, e.g. being specific or acting broadly, some data sets may fulfill the assumptions required by BackShift while others do not. Therefore, we considered only those results where the following conditions hold: (i) the joint diagonalization bootstrap confidence interval indicates that the joint diagonalization was successful; (ii) when refitting the model on subsamples of the data in the stability selection procedure, we require the joint diagonalization to be successful in at least 75 out of 100 runs. When these conditions were not met, we excluded the corresponding datasets from further consideration as the model assumptions of BackShift could be violated in these cases.

### Predictions

Figure 4 shows the causal relationships predicted by CLCD and BackShift. Predictions are aggregated over all different activators (for CLCD and BackShift) and inhibitors (for BackShift). For the subpopulations not shown for BackShift in Fig. 4, the graph returned by the stability selection procedure was either empty, or obtaining the point estimate itself was not successful (joint diagonalization did not succeed).

**Figure 4.**
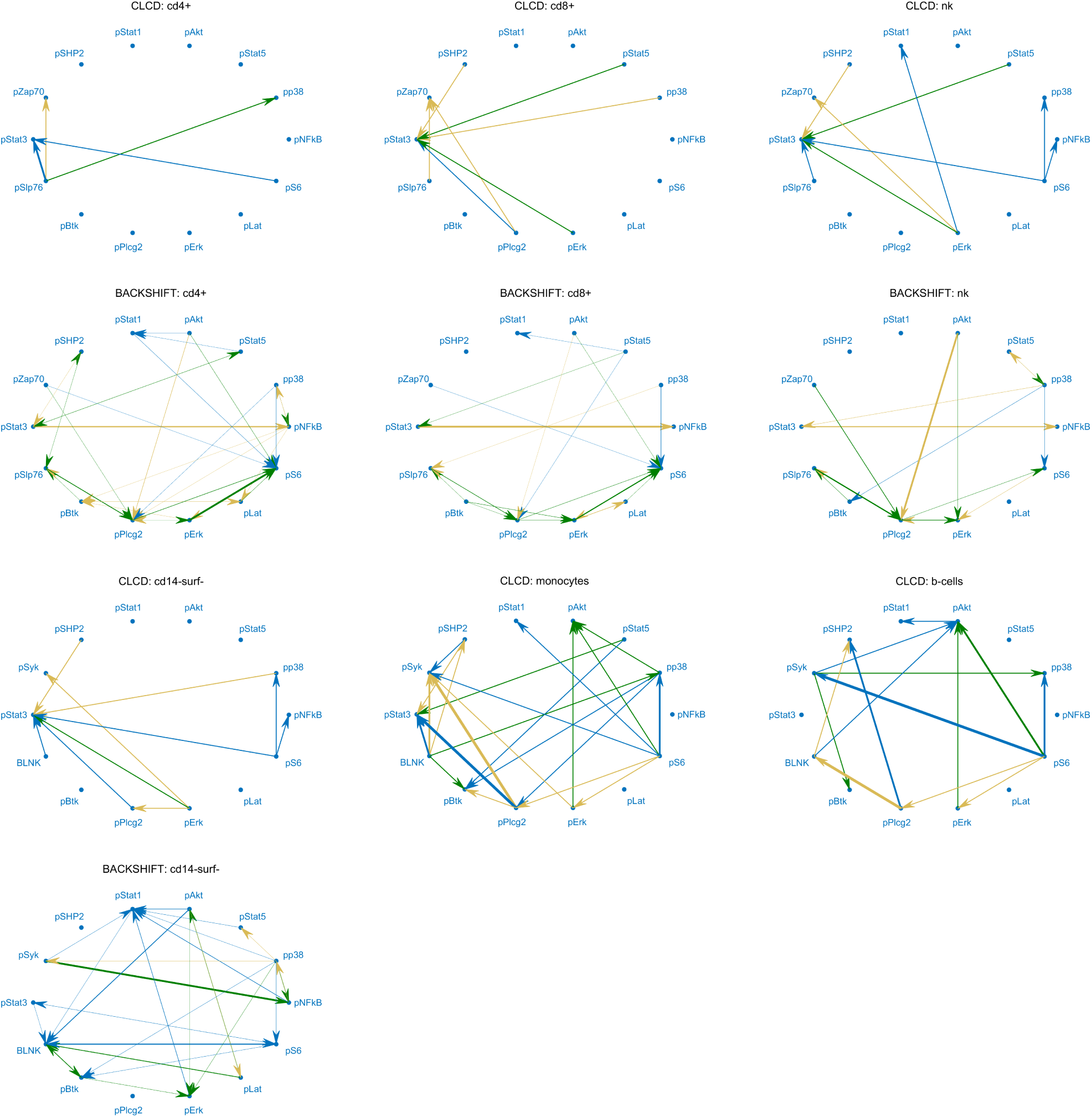
Predicted causal pairs for CLCD and backShift. CLCD outputs 82 different predictions (38 unique cause-effect pairs), while BackShift outputs 193 predictions (51 unique cause-effect pairs). BackShiftwas successful for 4 out of 14 populations: CD4+ and CD8+ T cells, Natural Killer (NK) cells, and CD14-surface- cells. CLCD results for the remaining subpopulations are grouped into CD14+surface- and monocytes (CD14+HLA-Dr-, CD14+HLA-Dr^high^, CD14+HLA-Dr^mid^, CD14-HLA-Dr-, CD14-HLA-Dr^high^, CD14-HLA-Dr^mid^) and B cells (IgM- and IgM+). Edge thickness corresponds to frequency of appearance in different contexts (activators, inhibitors, subpopulations where applicable). Green edges are confirmed in at least one KEGG pathway, while brown edges are found reversed in at least one KEGG pathway. A detailed list with the predictions can be found in Supplementary Tab. 3

### Reproducibility in independent data sets

Apart from the data used in the CLCD and BackShift pipelines, the same study^5^ includes time course data measuring protein response to several activators in PBMC samples across (a) eight different time points after activation (“time course data”), and (b) eight different donors (“8 donor data”), all measuring the same activators and proteins used in the main dataset with inhibitor treatments that was used above for CLCD and BackShift. Since BackShift demands different environments, it is not applicable in these additional data sets. In contrast, CLCD makes use of stimulated and unstimulated data, and the predictions should be consistent in matching conditions.

The exact CLCD pipelines are not applicable on the different time course and different donor data sets, so we used an (asymptotically equivalent) exact scoring scheme to test CLCD predictions, based on the BGEu score^18^. Baseline consistency was estimated using random predictions selected in a stratified manner, over 10 iterations.

Predictions are significantly consistent in all time points (Fig. 5, p-values: 0.03,10^−34^,10^−10^,10^−56^, 10^−34^,10^−52^,10^−9^,10^−7^ for 0,1,5,15,30,60,120 and 240 minutes, respectively). For the first time points after activation (0 & 1 minute), sparser networks where activation is not yet effective or has not yet reached the target protein are selected for the majority of the predictions (Supplementary Fig. 3). Activation is effective for most of the predictions after 5 minutes, and the same causal model (A→S→T) is confirmed for the majority of the predictions at 30 minutes. Activation effects decline after thirty minutes, returning to sparser selected networks. The majority of the predictions are not contradicted in the time course data.

**Figure 5.**
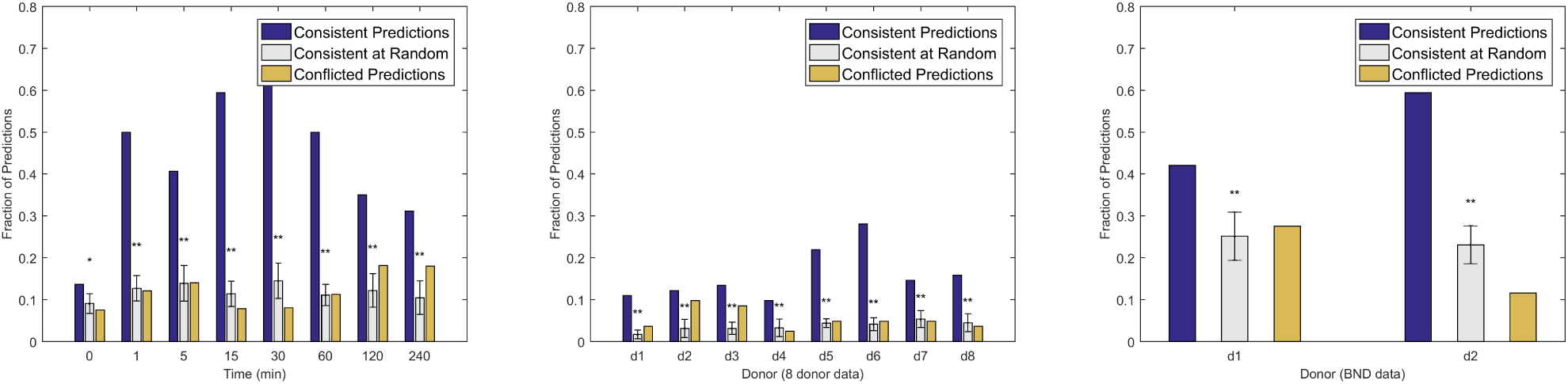
Reproducibility of the predictions in similar data sets. For every CLCD prediction (A ⟶ S ⟶ T), all possible networks were scored in an independent data set, and prediction is consistent if the same network is selected by the scoring scheme. In contrast, if the highest scoring network contradicts the predictions (i.e. includes a S ⟵ T edge), the prediction is considered conflicted. Predicted networks are found consistent in a variety of data sets, measuring the same variables and activators in (right) 8 distinct time points, (middle) 8 different healthy donors and (right) bone marrow data from two healthy donors in an independent study^19^. Asterisks denote statistical significance at 0.05(*) or 0.001(**). A one-tailed t-test is used to compare consistent predictions to consistency at random.

Similarly, the predictions are significantly consistent for all 8 healthy donors (Fig. 5, p-values: 10^−19^, 10^−5^, 10^−13^, 10^−4^, 10^−65^, 10^−54^, 10^−6^, 10^−8^). However, the actual percentage of consistent predictions is pretty low. A more careful inspection of the reproducibility results for the 8 donor data reveal that, in the majority of cases, protein levels are unaffected by the activation (see Supplementary Fig. 3), indicating possible problems in the experimental treatment.

To further test the reproducibility of CLCD predictions, we used not only the time course data and 8 donor data from the same study^5^ but in addition, we analyzed a completely independent dataset from another study measuring surface proteins and intracellular protein phosphorylation levels in bone marrow immune cells of two healthy human donors, 15 minutes after stimulation with various activators^19^ (denoted “bone marrow data”). Some of the proteins and activators are shared between the two studies (see Supplementary Tab. 1). While the studies are by no means identical, we tested whether similar phosphorylation patterns emerge in similar cell cell subpopulations upon activation with the same activators.

We used the same exact scoring scheme using the BGEu score^18^ to test CLCD predictions. The bone marrow data are gated according to a different gating hierarchy, resulting in subpopulations that do not have a one-to-one correspondence to the PBMC data. Each subpopulation specific triplet was therefore tested in all similar populations (B-cells, T-cells and NK cells, monocytes, dendritic cells). Despite differences in cell type, gating hierarchy and activation time, almost half of the predictions are consistent for each donor. (Fig. 5, donor 1: *p =* 10^−9^, donor 2: *p =* 10^−24^). Moreover, for networks that were not confirmed in the bone marrow data, the highest-scoring networks do not contradict the prediction (Supplementary Fig.3): for most inconsistent triplets, the full model is predicted, suggesting that the stimulation of the target protein could not be mediated by the target protein alone.

### Validation of causal predictions

Without targeted randomized control trials, causal claims are hard to prove. While the published data include responses of the measured proteins to increasing dosages of 27 inhibitors, the inhibitions are typically not absolutely target-specific, and definitely not surgical. One inhibitor that can be considered an exception is Rapamycin, which is a very specific inhibitor known to target the mTORc1 protein complex, while the closely related complex mTORc2 is largely unaffected. The mTORc1 kinase complex is directly upstream of the S6K protein, which phosphorylates S6 (to obtain pS6) if the complex is active^20^. Data from experiments using the inhibitor Rapamycin, thus, seem to be suitable to test causal predictions, for example it could be tested whether a predicted downstream target of pS6 would be affected by the presence of Rapamycin. Successful testing of such prediction requires the experimental system to show the expected behavior for known controls, that is, pS6 should decrease in the presence of Rapamycin. However, increasing doses of Rapamycin did not inhibit the phosphorylation of S6 in the public data we examined, as shown in the dose response curves for Rapamycin and its known target pS6, along with newly predicted targets of S6 in response to Rapamyin in Supplementary Fig. 5 and 6.

**Figure 6.**
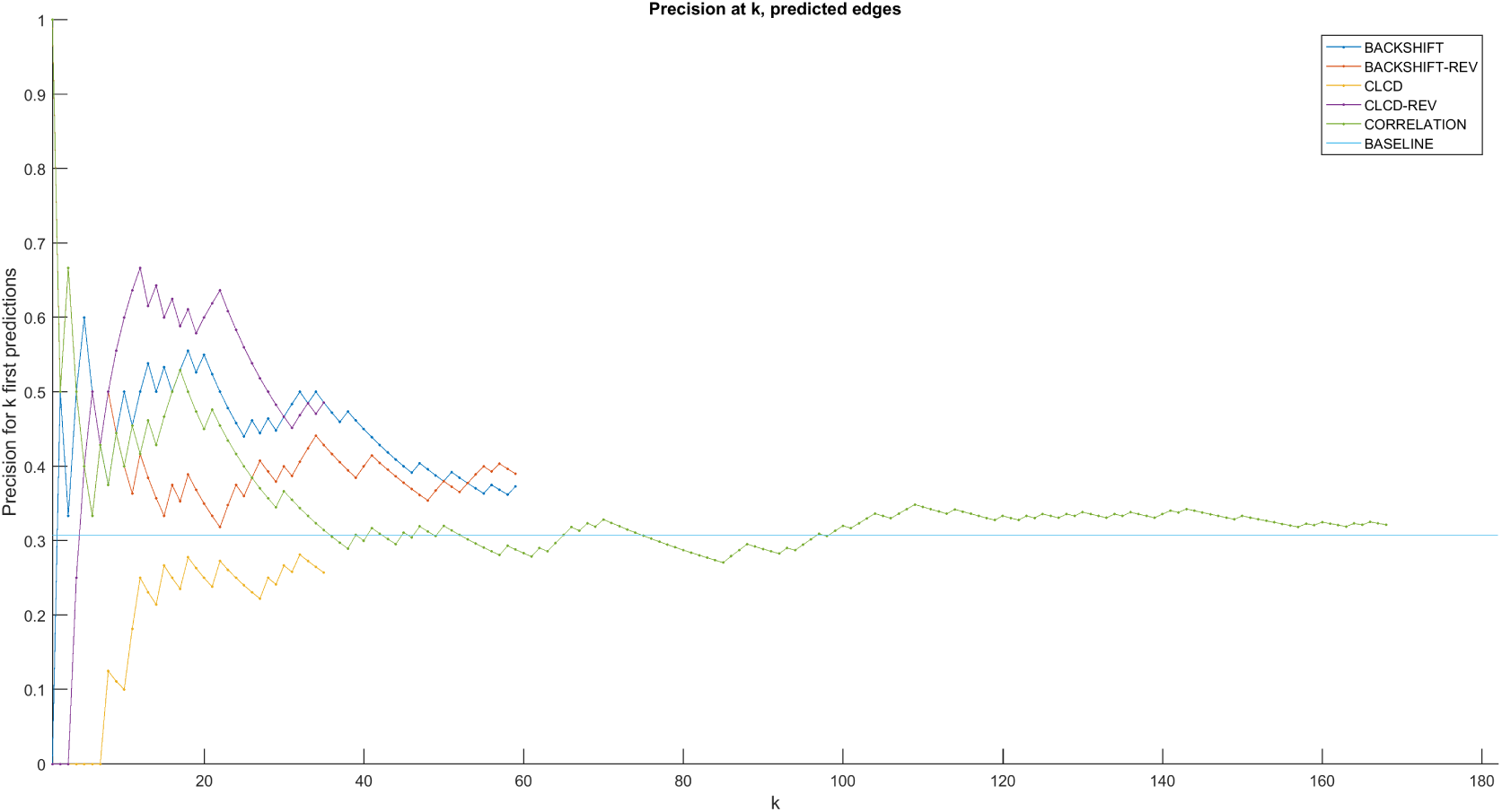
Precision of CLCD and BackShift predictions in KEGG human pathways. Cause-effect predictions were ranked according to frequency of appearance and then tested in KEGG human pathways. Out of 182 possible pairs, 56 pairs have a (direct or indirect) causal relationship in KEGG (baseline prediction).BackShiftperformed better than CLCD and mere correlation, but no method was significantly better than chance.

This indicates that either the cells in the experimental source data do not react as expected to respective stimulators and/or inhibitors, the chosen time points were not optimal for the respective readout, or the antibodies to detect the respective phosphoproteins did not work. The latter can be excluded however for pS6 since an inhibition of pS6 signal is seen in CD4+ T cells with the (unspecific) pan kinase inhibitor Staurosporine (see original publication^5^ Fig. 4d). As also displayed in the original publication, it can be furthermore concluded that although CD4+ and CD8+ T cells reacted with S6 phosphorylation to PMA/Iono stimulation, no inhibition of this pS6 signal was seen when the S6 inhibitor Rapamycin was applied (see original publication^5^ Supplemental Fig. S19).

In another attempt to validate the predictions, we consulted the KEGG database which includes a collection of human protein signaling pathways based on literature from a multitude of different experimental systems. While the list of causal relationships documented in the pathways is not exhaustive, we examined the status of predicted relationships in the KEGG pathways. The 301 KEGG pathway entries corresponding to human pathways were downloaded and causal ancestry relationships among the 14 measured intracellular proteins were automatically mined using KEGGParser^21^.

Figure 6 shows the precision of ranked predictions in the KEGG validation. BackShift performs slightly better than correlation, however none of the two methods was significantly better than random. Nine out of 38 CLCD unique predictions (regardless of activator and subpopulation) were identified in KEGG. Intriguingly, for 17 out of 38 predictions, the reverse relationship was identified in KEGG. BackShift achieved better performance: 22 out of 60 predictions were confirmed in KEGG pathways, while 24 out 60 predictions were found reversed. However, notice that this comparison is not very precise: the predictions were tested in *all* KEGG human pathways, regardless of the subpopulation and activator used to stimulate the cells. Since different cells can react with different pathways and to diverse stimulators based on the receptors, signaling components and regulators they express, this may contribute to mismatch of the pathway relationships even though some may still be true in reality.

## Discussion

Biological data are often noisy, and highly dependent on the quality and availability of the used tools. In the analyzed data, a limitation of the CyTOF method is that the signal can only be as good as the available antibodies to detect the respective proteins, and the utilized inhibitors are not all absolutely specific for their desired target. Furthermore, the use of human primary cells naturally leads to donor variability with regard to response to stimulation, inhibition and expression of the measured proteins, as compared to for example measurements on more uniform cell lines. Another factor that complicates causal analyses is the frequent presence of (negative and positive) feedback loops in biological signaling systems.

Nevertheless, mass cytometry data have several characteristics that make them suitable for causal discovery: namely, they do not suffer from population averaging and come in adequately large sample sizes. Flow cytometry data, which are very similar in principle, have been used successfully in the past^4^.

In this work, we examined the performance of two causal discovery methods on a collection of mass cytometry data. The methods are based on fundamental causal principles, and use multiple data sets and/or different experimental conditions to increase robustness. However, we found that even basic methods do not have perfect reconstruction accuracy, and often disagree with background knowledge (such as the KEGG pathways). On the other hand, results are shown to be reproducible in independent data sets, showing statistical patterns associated with certain causal models are consistent across different studies.

For automatic causal discovery, these results illustrate the need for (a) systematic study of the effect of violations of causal assumptions to existing algorithms, and (b) novel causal discovery methods that focus on conservative, quantitative, testable predictions in the presence of confounders and feedback cycles. Moreover, these results indicate the importance of practical applications for causal discovery algorithms; such interdisciplinary applications must be carefully selected, designed and tested in collaboration with domain experts, who can also guide the evaluation and extension of algorithms based on their domain-specific knowledge.

## Methods

Gated data from^5^ and^19^ were downloaded from Cytobank (http://cytobank.org/). For each protein pair (*P*_1_, *P*_2_) and Activator (*A*), CLCD performs 6 tests of independence:

- *P*_1_ ╨*A*|Ø, *P*_2_ ╨*A*|Ø: t-test.
- *P*_1_ ╨*P*_2_|Ø, *P*_1_ ╨*P*_2_|*A*: Fisher z-test^22^.
- *P*_1_╨*A*|*P*_2_, *P*_2_ ╨*A*|*P*_1_: logistic test^23^.

For BackShift, the causal system over *p* measurement variables **x** is defined by the set of linear relationships

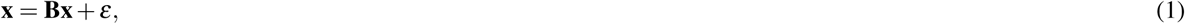
where **B**_*i, j*_ is non-zero if *x_j_* is a cause of *x_i_*, and the error term ε is a p-dimensional random variable with mean 0 and positive semi-definite covariance matrix Σ_ε_.

The connectivity matrix **B** fully describes the causal structure of the measured variables: the non-zero coefficients of **B** correspond to the direct causal relationships, and the value of each coefficient describes the strength of the corresponding relationship. Allowing Σ_ε_ to be non-diagonal allows for possible latent confounders, and restricting the diagonal elements of **B** to be zero forbids self-loops. This is necessary for model identifiability.

For estimating the connectivity matrix **B**, BackShift relies on observations of the system under shift interventions in different environments *e* ∈ *ℰ*

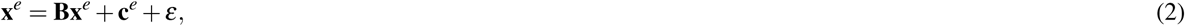
where **c***^e^* ∈ ℝ^*p*^ are uncorrelated interventions (the covariance matrix of **c***^e^* is a diagonal matrix). Thus, the intervention is allowed to perturb each variable, while the causal relationships between the variables remain the same. Identifying matrix **B** is equivalent to fully identifying both the causal structure (non-zero elements) and the strength of causal relationships.

Estimation of **B** is based on the relation between the covariance matrices (i) 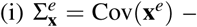 – the covariance of **x** in environment *e* (observed), (ii) 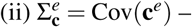 – the covariance of **c** in environment *e* (unobserved) and (iii) 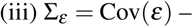 – the covariance of the noise (unobserved).

Exploiting the structure of the model, BackShift finds **B** ∈ ℝ*^p×p^* such that ∀*e* ∈ *ℰ*

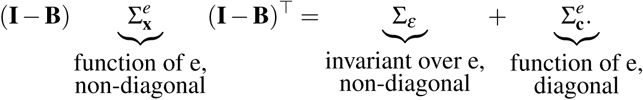

The estimation relies on a simple joint matrix diagonalization, applied to differences between covariance matrices, and a reformulation of the linear sum assignment problem. This amounts to a computational complexity of *O(np^2^)* where *n* is the total number of observations across all environments. For further details about the method, we refer to^16^. Failure of the joint diagonalization algorithm indicates some model violation. The success or failure of the joint diagonalization algorithm is assessed by a parametric bootstrap approach.

For the stability selection, the parameters were chosen as 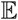(*V*) = 5,*n_sim_=100* and *π_thr_ =* 0.75. This implies that the estimated network only contains those edges that were found more than 75 times when refitting the model on 100 random subsamples of the data. Thus, stability selection allows us to assess whether BackShift yields consistent estimates when different subsamples of the same dataset are used. Moreover, only these stable results are included in the final output. In the analysis, the expected number *V* of falsely selected variables was set to 5. In general, decreasing 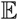(*V*) leads to sparser graphs.

## Acknowledgements

The authors would like to thank Karen Sachs for offering valuable insights on mass cytometry, and Joris Mooij for helpful discussions. ST, VL and IT were supported by the European Research Council under the European Union’s Seventh Framework Programme (FP/2007-2013) / ERC Grant Agreement n. 617393.

## Author contributions statement

S.T., V.L. and I.T. conceived the idea. S.T. and V.L. preprocessed the data. S.T., V.L. and C.H.D. analyzed the data. S.T., V.L. and A.S designed the validation analysis. All authors reviewed the manuscript.

